# ABA represses TOR and root meristem activity through nuclear exit of the SnRK1 kinase

**DOI:** 10.1101/2021.12.27.474243

**Authors:** Borja Belda-Palazón, Mónica Costa, Tom Beeckman, Filip Rolland, Elena Baena-González

## Abstract

The phytohormone abscisic acid (ABA) promotes plant tolerance to major stresses like drought, partly by modulating plant growth. However, the underlying mechanisms are poorly understood. Here, we show that cell proliferation in the *Arabidopsis thaliana* root meristem is controlled by the interplay between three kinases, SNF1-RELATED KINASE 2 (SnRK2), the main driver of ABA signaling, the SnRK1 energy sensor, and the growth-promoting TARGET OF RAPAMYCIN (TOR) kinase. Under favorable conditions, the SnRK1α1 catalytic subunit is enriched in the nuclei of root cells and this is accompanied by normal cell proliferation and meristem size. Depletion of SnRK2s in a *snrk2*.*2 snrk2*.*3* double mutant causes constitutive cytoplasmic localization of SnRK1α1 and a reduction in meristem size, suggesting that, under non-stress conditions, SnRK2s enable growth by retaining SnRK1α1 in the nucleus. In response to elevated ABA levels, SnRK1α1 translocates to the cytoplasm and this is accompanied by inhibition of TOR, decreased cell proliferation and meristem size. Blocking nuclear export with leptomycin B abrogates ABA-driven SnRK1α1 relocalization to the cytoplasm and the inhibition of TOR. Fusion of SnRK1α1 to an SV40 nuclear localization signal leads to defective TOR repression in response to ABA, demonstrating that SnRK1α1 nuclear exit is a premise for this repression. Altogether, we demonstrate that SnRK2-dependent changes in SnRK1α1 subcellular localization are crucial for the regulation of TOR activity and root growth in response to ABA. Such swift relocalization of key regulators may represent a more general strategy of sessile organisms like plants to rapidly respond to environmental changes.

## Main Text

The phytohormone abscisic acid (ABA) plays major roles in plant stress responses. ABA signals are transduced through a well-established pathway whose main effectors in *Arabidopsis* are SNF1-RELATED PROTEIN KINASE 2.2 (SnRK2.2), SnRK2.3, and SnRK2.6 (1). ABA promotes plant adaptation partly by modifying developmental programs, having a major impact *e*.*g*. on root architecture (2). ABA modulates primary root (PR) and lateral root (LR) growth through interactions with other hormones, ultimately affecting cell division and elongation by mechanisms still poorly understood (2). We recently uncovered an intimate connection between ABA and SnRK1 signalling that is crucial for shaping root architecture in a TARGET OF RAPAMYCIN (TOR)-dependent manner (3). TOR is a protein kinase complex that promotes cell proliferation, with its inactivation causing reduced root meristem size and defective PR growth (4). SnRK1 is an evolutionarily conserved protein kinase complex that is activated when energy levels decline during stress, conferring protection partly by limiting growth (5). SnRK1 is also activated by ABA, enabling plants to repress growth when water is scarce (3). Under favorable conditions, the main catalytic subunit SnRK1α1 is sequestered by SnRK2-containing repressor complexes, allowing TOR to be active. In response to ABA, these complexes dissociate, releasing SnRK2 and SnRK1α1, which inhibits TOR and growth (3). Consistent with this model, the *snrk2*.*2 snrk2*.*3* double mutant (*snrk2d*) shows markedly defective PR growth under favorable conditions due to aberrant repression of TOR activity (3). This defect is fully rescued by the *snrk1α1* mutation, demonstrating its SnRK1α1-dependency (3).

To investigate how TOR and root growth are controlled by SnRK2 and SnRK1, we examined the root meristems of Col-0 control seedlings, the *snrk2d* mutant, and its cross with *snrk1α1* (*snrk2d/1α1*) (Fig. 1*A*). As previously reported (6, 7), ABA treatment reduced the number of root meristematic cells, leading to smaller meristems in Col-0. In contrast, *snrk2d* showed a reduction in meristem size and cell number already in control conditions (mock treatment), and this was fully rescued by the *snrk1α1* mutation. Consistent with the ABA hyposensitivity of *snrk2d* and *snrk2d/1α1* plants (3), the addition of ABA did not further decrease meristem size or cell number in these mutants (Fig. 1*A*). These meristem phenotypes correlate well with the PR length previously observed in these mutants and conditions (3) and suggest that reduced cell proliferation contributes to the reduced PR length of *snrk2d* in control conditions, mimicking the situation of Col-0 plants upon ABA treatment.

**Figure 1.**
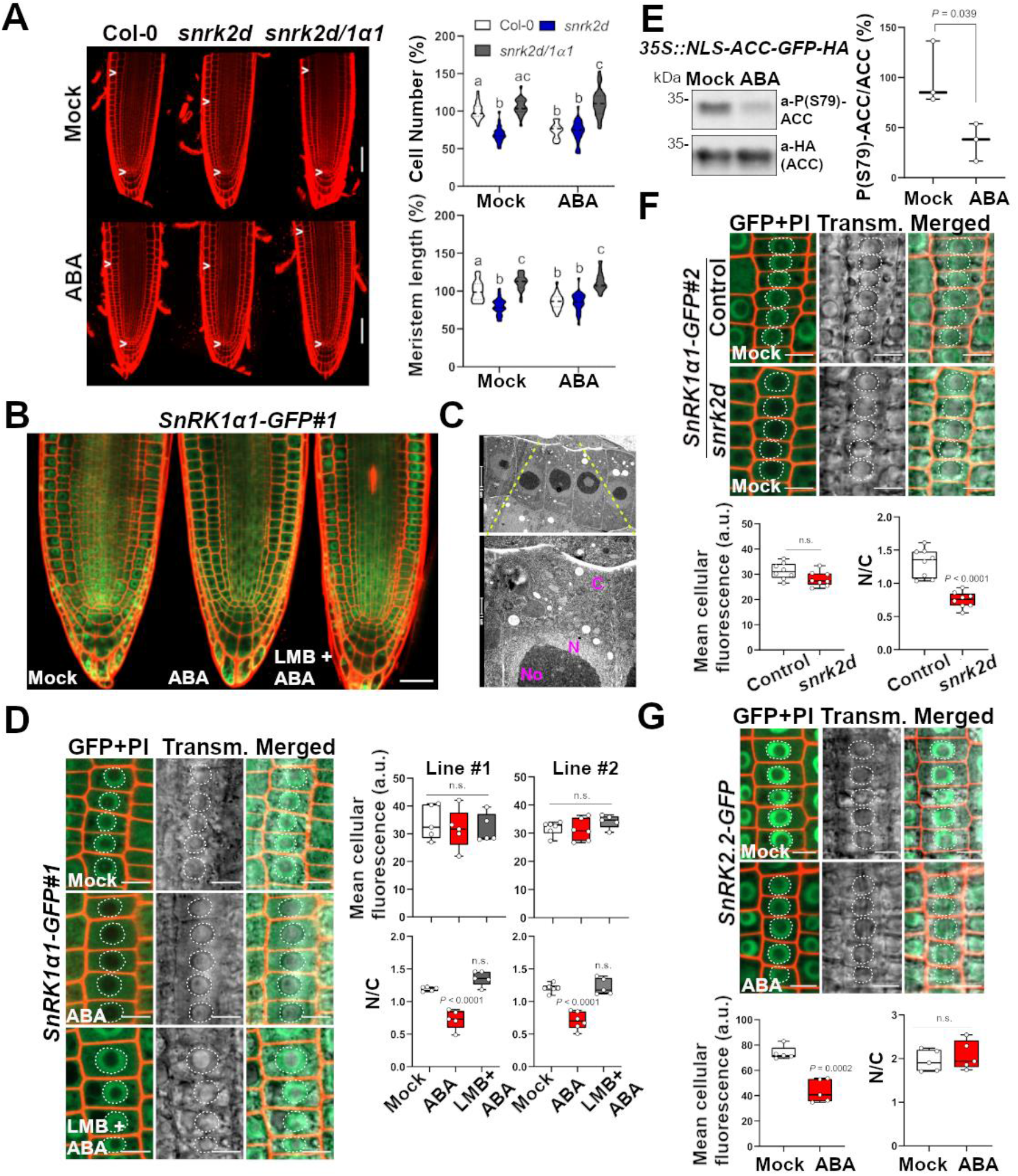
Impact of ABA and SnRK2s on SnRK1α1 subcellular localization and cell proliferation in the root apical meristem. (*A*) Meristems of Col-0, *snrk2d* and *snrk2d/1α1* 7d-old seedlings with or without ABA treatment (50 µM, 48h). Arrowheads: region for cortical cell number and meristem length quantifications (violin plots). *n*=23-24; *p* < 0.05, one-way ANOVA with Tukey ‘s HSD test. Scale bar: 50 µm. (*B*) Root apices of 4d-old *SnRK1α1-GFP#1* seedlings with or without ABA treatment (50 µM, 3h). Scale bar: 30 µm. (*C*) Electron micrograph of meristematic epidermal cells with magnification (*lower panel*) showing the cellular ultrastructure. C: cytoplasm; N: nucleus; No: nucleolus. Scale bars: upper, 5 µm; lower, 1 µm. (*D*) Left panels, SnRK1α1-GFP subcellular localization in epidermal cells of 4d-old root meristems. Scale bar: 10 µm. Dotted lines: nuclear boundary. Right panels, quantification of SnRK1α1-GFP mean cellular fluorescence and nucleus- to-cytosol (N/C) ratios. *n*=5-6; one-way ANOVA with Dunett’s test. (*E*) Nuclear SnRK1 activity in 8d-old roots of the *NLS-ACC* reporter line with or without ABA treatment (50 µM, 3h). Left panel, representative immunoblot. Right panel, quantification of ACC phosphorylation levels (P(S79)-ACC/total ACC). *n*=3, two-tailed Student t-test. (*F*) Impact of the presence (control, *SnRK1α1-GFP#2)* or absence (*SnRK1α1-GFP#2; snrk2d*) of SnRK2s on SnRK1α1-GFP localization assessed as in (*D*). *n*=8; two-tailed Student t-test. (*G*) Expression and subcellular localization of SnRK2.2-GFP in epidermal cells of 4d-old root meristems treated and quantified as in (*D*). *n*=5; two-tailed Student t-test. *PI*, propidium iodide. *ns*, non-significant.

We next hypothesized that regulation of SnRK1 and growth by ABA and SnRK2s could involve changes in SnRK1α1 subcellular localization. Firstly, the subcellular localization of SnRK1α1 is central to its function (8). Secondly, SnRK1 and SnRK2 are highly enriched in the nuclei of root cells (3). Thirdly, SnRK2-harboring SnRK1 repressor complexes localize to the nucleus in *Nicotiana benthamiana* epidermal cells (3). Fourthly, *in planta*, the TOR complex subunit RAPTOR1B interacts with SnRK1α1 (3, 9) in the cytosol (10). We therefore monitored SnRK1α1 subcellular localization in mock- or ABA-treated seedlings in two independent lines expressing SnRK1α1-GFP from its own regulatory regions (*SnRK1α1-GFP#1* and *SnRK1α1-GFP#2*). In control conditions, SnRK1α1 displayed a known nuclear and cytoplasmic localization (11), with the latter exhibiting a ring-shaped pattern characteristic of nuclear proteins that are absent from the nucleolus in meristematic cells (12) (Fig. 1*B*). Electron micrographs confirmed that a ring-shaped nuclear region surrounds a large-sized nucleolus in these cells (Fig. 1*C*). Upon ABA treatment, the nuclear signal appeared to decline (Fig. 1*B*). Quantification of the GFP signal revealed overall similar SnRK1α1-GFP levels in mock- and ABA-treated roots but a reduction in the nucleus to cytoplasm ratio (N/C) from 1.19 (line #1) and 1.2 (line #2) in mock to 0.73 (line #1) and 0.71 (line #2) in ABA (Fig. 1*D*), suggesting that ABA induces SnRK1α1 relocalization from the nucleus to the cytoplasm. Accordingly, the ABA effect was fully abolished by the nuclear export inhibitor Leptomycin B (LMB; N/C ratio=1.36 and 1.23 in lines #1 and #2; Fig. 1*D*). To investigate the potential consequences of SnRK1α1 nuclear exit for SnRK1 activity, we used a line expressing a rat ACETYL-COA CARBOXYLASE (ACC) peptide, a well-established target of the SnRK1 mammalian ortholog, fused to an SV40 nuclear localization sequence (NLS) [*35S::NLS-ratACC-GFP-HA*, referred to as *NLS-ACC;* (13)]. Phosphorylation of the synthetic NLS-ACC protein serves as readout of SnRK1 nuclear activity (13). Treatment of the *NLS-ACC* seedlings with ABA led to a clear reduction in ACC phosphorylation as compared to mock (Fig. 1*E*). This indicates a decline in nuclear SnRK1 activity that is consistent with the observed nuclear exit of SnRK1α1 in response to ABA (Fig. 1*B, D*).

Given the marked reduction in meristem size (Fig. 1*A*) and PR length (3) of the *snrk2d* mutant, we investigated the role of SnRK2s in SnRK1α1 localization using the cross of *snrk2d* with the *SnRK1α1-GFP#2* line (Fig. 1*F*). In mock conditions, SnRK1α1-GFP was barely present in the nuclei of *snrk2d* cells, with most of the GFP signal being cytoplasmatic and yielding a N/C ratio of 0.75 (Fig. 1*F*), comparable to the ABA-treated *SnRK1α1-GFP#2* control (Fig. 1*D*). SnRK2.2-GFP suffered from overall protein degradation, but there was no change in its subcellular localization (Fig. 1*G*). Altogether, these results show that ABA triggers the translocation of SnRK1α1 to the cytoplasm and that SnRK2s are required for retaining SnRK1α1 in the nucleus when ABA is not present. Intriguingly, low energy stress leads to an increase in nuclear SnRK1 activity (13), reinforcing the view that different signals activate different SnRK1 complexes and that SnRK2 kinases are only required for the activation of SnRK1 in response to ABA (3).

Given that SnRK1α1 is required for the inhibitory effect of ABA on TOR and growth (3), we investigated whether nuclear exit of SnRK1α1 is a premise for such inhibition, using the phosphorylation of ribosomal protein S6 (RPS6^S240^) as readout of TOR activity (3). While LMB treatment had no significant impact (Fig. 2*A*, left panels), LMB was able to block the repression of RPS6 phosphorylation triggered by ABA (Fig. 2*A*, right panels). Moreover, as compared to control plants expressing wild-type SnRK1α1 (*control-α1*), the repression of TOR by ABA was also defective when SnRK1α1 was fused to an SV40 nuclear localization sequence (NLS) that favors its presence in the nucleus [*NLS-α1*; (8); Fig. 2*B*]. This demonstrates that SnRK1α1 nuclear exit is necessary for inhibiting TOR in response to ABA. Furthermore, the fact that nuclear export is crucial for repressing TOR activity (Fig. 2*A*) but only SnRK1α1 (Fig. 1*D*), and not SnRK2.2 (Fig. 1*F*), translocates to the cytoplasm in response to the hormone, suggests that the previously reported role of SnRK2s in this process (3, 14) is indirect *via* SnRK1α1 regulation.

**Figure 2.**
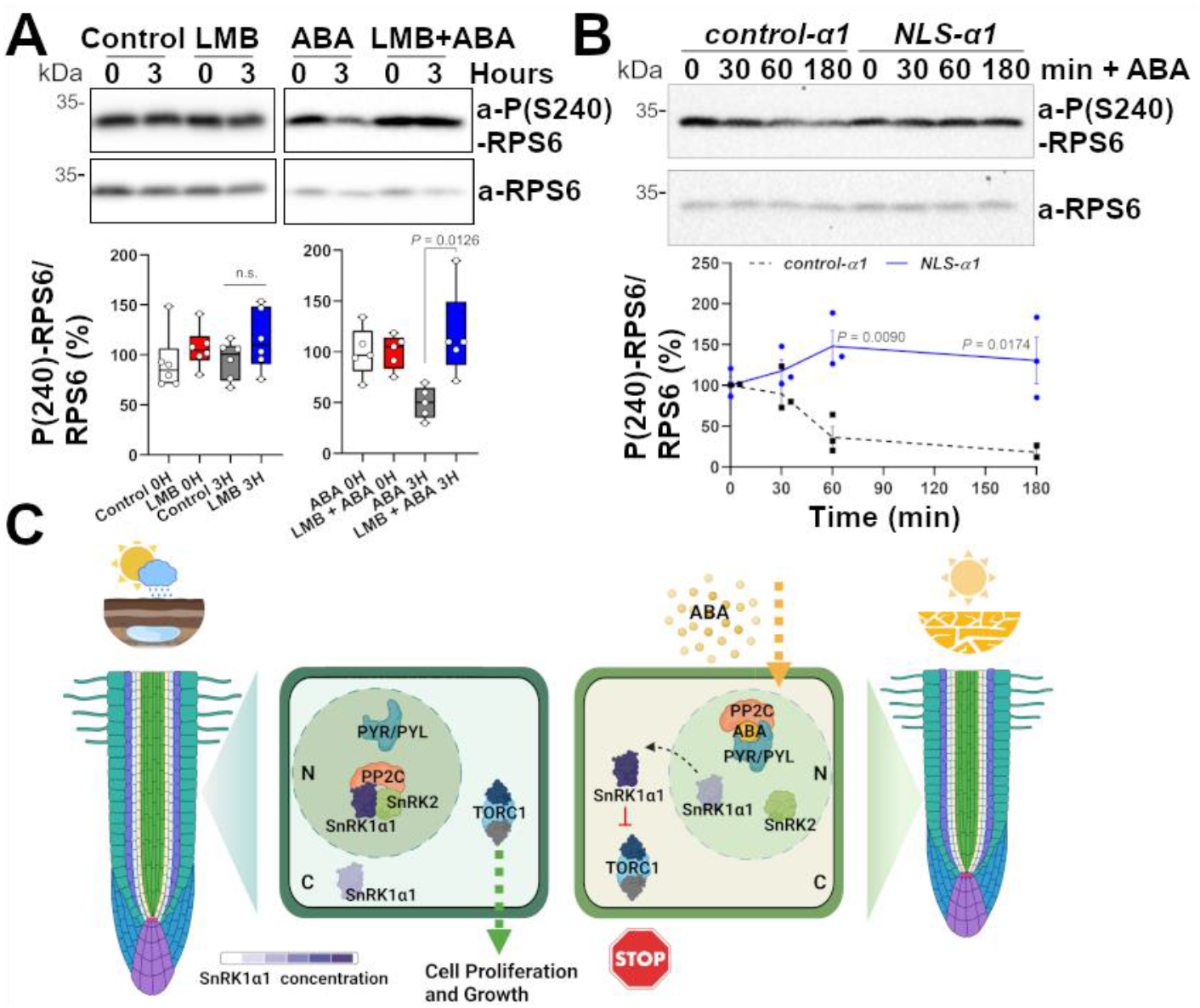
Impact of SnRK1α1 subcellular localization on TOR signaling and root growth. (*A*-*B*) Upper panels, representative immunoblots of RPS6^S240^ phosphorylation (phospho-RPS6/total-RPS6) in Col-0 (*A*) or *control-α1* and *NLS-α1* (*B*) seedlings treated with or without ABA (50 µM, 3h) or LMB (2.5 µM, 3h or 1h prior to ABA addition). Images from the same gel were cropped for showing *α1* and *NLS-α1* contiguously. Lower panels, quantification of phospho-RPS6/total-RPS6 from immunoblots. (*A*) *n*=5-6; (*B*) *n*= 3; error bars, SEM; two-tailed Student *t*-test. (*C*) Under favorable conditions, SnRK1α1 is sequestered in the nucleus by repressor complexes containing SnRK2 [and a PP2C (3)], enabling TOR activity in the cytoplasm, cell proliferation and root growth. Dissociation of these complexes in ABA by the hormone-bound PYR/PYL/RCAR receptors releases SnRK1α1 which exits the nucleus and inhibits TOR and growth. *TORC1*, TOR Complex 1; *N*, nucleus; *C*, cytoplasm. Created with BioRender.com.

We conclude that root growth is modulated by ABA through changes in SnRK1α1 subcellular localization, allowing control of TOR activity and cell proliferation in the root meristem in accordance to *e*.*g*. water availability (Fig. 2*C*). When conditions are favorable, SnRK1α1 is sequestered in the nucleus by SnRK2-containing repressor complexes. Dissociation of these complexes in response to ABA releases SnRK1α1, which translocates to the cytoplasm and inhibits TOR activity and growth. Mechanistic understanding of how growth is repressed by ABA is highly relevant, as it may provide new means to manipulate growth-defense trade-offs in plants, enhancing stress tolerance without compromising growth and productivity.

## Materials and Methods

Experimental details are provided in SI Appendix.

All study data are included in the paper.

## Acknowledgments

We thank J-K. Zhu, M. Bennett and C. Koncz for *snrk2d, SnRK2*.*2-GFP*, and *SnRK1α1-GFP#1* seeds. We also thank the IGC Facilities for excellent plant care (*Plant*), sample processing and imaging (*Electron Microscopy*) and technical advice (*Advanced Imaging*; supported by PPBI-POCI-01-0145-FEDER-022122). This work was funded by FCT [UIDB/04551/2020, LISBOA-01-0145-FEDER-028128, PTDC/BIA-BID/32347/2017], and by EU Horizon 2020 programme (BBP; H2020-WF-2018-2020/H2020-WF-01-2018, Grant 867426).

## Notes

### Competing Interest Statement

The authors have declared no competing interest.

### Summary of Updates

In addition to our confocal microscopy analyses, we now show that ABA treatment leads to a decrease in the nuclear activity of SnRK1 (Figure 1E of revised MS), supporting the conclusion that ABA triggers the exit of SnRK1 from the nucleus in whole roots. This evidence relies on a recently published Arabidopsis reporter line expressing a synthetic protein (NLS-ratACC-GFP-HA; Muralidhara et al., 2021 PNAS), whose phosphorylation is a direct read-out of SnRK1 nuclear activity and can be detected by immunoblotting with commercial antibodies. We also used the NLS-ACC reporter line to analyze SnRK1 nuclear activities in dissected root tips and upper root regions. These analyses left no doubt that the decline in SnRK1 nuclear activity in response to ABA occurs similarly in the differentiated regions of the root (data not shown).

