## Extended methods for "ABA represses TOR and root meristem activity through nuclear exit of the SnRK1 kinase"

### Plant material and growth

All *Arabidopsis* plants used in this study are in the Columbia (Col-0) background. Unless otherwise specified, plants were grown under long-day conditions (16 h light, 100 µmol $m^{-2}$ $s^{-1}$, 22 °C/8 h dark, 18 °C) on 0.5X MS medium (0.05% MES and 0.8% phytoagar).

All the following lines were previously described: *snrk2d* [*snrk2.2* (GABI-Kat 807G04)/ *snrk2.3* (SALK_107315); (1)], *snrk2d/1α1* [*snrk1α1-3* (GABI_579E09); (2)], *SnRK2.2-GFP* #2.2 (3), *SnRK1α1-GFP#1* (4), *SnRK1α1-GFP#2* (5), *control-α1* and *NLS-α1* (6), *NLS-ACC* (7). The *SnRK1α1-GFP#2 snrk2d* transgenic line was obtained by crossing *SnRK1α1-GFP#2* to *snrk2d*.

### Analyses of root apical meristem development

For assaying root apical meristem development, seedlings were grown vertically for 5 days in 0.5X MS and transferred to 0.5X MS plates with or without 50 µM ABA where seedlings were allowed to grow vertically for 2 more days. Roots were stained for 2 min with an aqueous solution of propidium iodide (PI; 10 µg/mL) and images were acquired on a Zeiss LSM980 system [using Airyscan SR/Multiplex4Y mode] equipped with two PMT and one GaAsP, using a 40× 1.1NA water immersion objective. For the visualization of cell walls, pinholes were adjusted to 1 Air Unit (561 nm/600-660 nm). Post-acquisition image processing was performed using Zeiss's ZEN Blue v3.0 imaging software and ImageJ (http://rsb.info.gov/ij/). Quantification of cortical cell number and meristem size was done from the region comprised between the quiescent center and the beginning of the transition zone (defined as the point where cortical cell length is first doubled). Meristem length was measured at the center of the vascular bundle. Measurements were performed from 23–24 seedlings per genotype and condition and grown as 3 independent batches.

### Subcellular localization analyses by CLSM

The localization of SnRK1α1 and SnRK2.2 was investigated in roots of *SnRK1α1-GFP#1*, *SnRK1α1-GFP#2, SnRK1α1-GFP#2 snrk2d* and *SnRK2.2-GFP* seedlings grown vertically on 0.5× MS plates for 4 days (primary roots) or 9 days (lateral roots). On day 4, seedlings were transferred to liquid 0.5× MS one hour after the onset of the lights and allowed to acclimate for two hours. Seedlings were thereafter treated with mock, 50 µM ABA or 50 µM ABA + 2.5 µM LMB for 3h. In the case of the combined treatment of LMB and ABA, seedlings were preincubated with LMB for one hour during the acclimation period before the addition of ABA. Roots were stained with PI and images were acquired on a Zeiss LSM980 system as described above for the visualization of the cell walls. For the visualization of GFP, pinholes were adjusted to 1 Air Unit (488 nm/500-530 nm). For quantitative analysis of GFP, the power of the 488 nm laser was set at 3.0% transmission to gain master of 800. Post-acquisition image processing was performed using with Zeiss's ZEN Blue v3.0 imaging software and ImageJ (http://rsb.info.gov/ij/).

The nuclear/cytoplasmic localization of SnRK1α1 and SnRK2.2 was assessed using CLSM and ImageJ software, calculating the N/C ratio (N/C = Mean Nuclear Fluorescence Intensity / Mean Cytoplasmic Fluorescence Intensity, with Mean Fluorescence Intensity being the ratio between the total fluorescence intensity measured and the number of pixels measured) (8). The area of the nucleus was determined by analyzing the bright field acquired through the transmitted light mode whilst the area of the cytoplasm was selected using the PI signal as a reference. Quantification of mean cellular fluorescence and nucleus-to-cytosol ratios was done from 5 root tips, each consisting of the average of 5 meristematic epidermal cells.

**Transmission electron microscopy**

Root tip samples were vitrified in 10% BSA with 8% methanol using a Wohlwend Compact 2 High Pressure Freezer (Engineering Office M. Wohlwend GmbH) prior to processing using an AFS2 with the FSP robot (Leica Microsystems) in a solution of 0.25% glutaraldehyde, 0.1% UA in dry acetone for 48 hours at -80°C. Samples were then warmed up to -50°C at a rate of 1°C/hr. The fixative was washed out with acetone three times for 10 minutes each and then samples were infiltrated in an increasing concentration of Lowicryl HM20 (22%, 33%, 66%, 100% x 3) prior to UV polymerization. Sections of 70nm were cut using an ultra45 diamond knife (Diatome) on a UC7 Ultramicrotome (Leica Microsystems) and collected on slot grids coated with 1% formvar in chloroform. The sections were post-stained sequentially with uranyl acetate and lead citrate for 5 minutes each and then imaged on a Tecnai G2 Spirit BioTWIN Transmission Electron Microscope (TEM) from FEI operating at 120 keV and equipped with an Olympus-SIS Veleta CCD Camera.

***In planta* SnRK1 activity assay**

Arabidopsis *NLS-ACC* seedlings were grown vertically for 8 days on calibrated Nytex mesh (pore size 30 µm) on solid medium (0.5X MS). On day 9, two hours after the onset of the lights, the mesh squares holding the seedlings were transferred to new solid medium plates with or without 50 µM ABA and returned to the growth chamber for 3h. Root tissues were thereafter separated from aerial parts and ground to a fine powder in liquid nitrogen. Root tissue powder (30 mg) was mixed with 2x Laemmli buffer and incubated on ice for 20 min, vortexing every 5 min. Samples were thereafter boiled for 10 min, cooled on ice for 5 min, and centrifuged at 12000g for 5 min at 4°C to clear homogenates. The resulting supernatants (15 µL) were analyzed by Western Blot with anti-P(S79)-ACC (1:1000, 1673661S, Cell Signaling) and anti-HA-HRP (1:2000, 12013819001, Roche) antibodies.

**ABA time course and RPS6$^{S240}$ phosphorylation assays**

The ability of the indicated genotypes to repress RPS6$^{S240}$ phosphorylation in response to ABA was analyzed as described (2). Seedlings were grown vertically for 6 days on solid medium (0.5× MS +

0.5% sucrose) and were thereafter transferred to liquid medium (0.5× MS medium + 0.5% sucrose) in 6-well tissue culture plates (10 seedlings per 9.5 $cm^2$ well containing 1 mL of medium) where they grew for the following 6 days. The liquid medium was refreshed 8h before the beginning of the last night and on the following day, samples were collected 2h after the onset of the lights (T0). Remaining seedlings were then treated with 50 µM ABA and/or 2.5 µM LMB for the indicated periods of time. In the case of the combined treatment of LMB and ABA, seedlings were preincubated with 2.5 µM LMB for 1h before the addition of ABA. Following the indicated treatments, samples were ground to a fine powder in liquid nitrogen and immediately placed in extraction buffer [50 mM Tris-HCl pH 8.0, 150 mM NaCl, 0.1% NP-40, 3 mM DTT, 50 µM MG-132, Phosphatase Inhibitor Cocktails 2 and 3 (Sigma; 20 µL each per 10 mL of extraction buffer) and cOmplete™ Protease Inhibitor Cocktail (Roche, 1 tablet per 10 mL of extraction buffer)] for total protein extraction (150 µL of buffer per 100 mg of ground tissue). Homogenates were cleared by centrifugation at 12000 g for 15 minutes at 4°C and supernatants were recovered for subsequent analyses. 50 µg of total protein extract of each sample were analyzed by Western Blot with anti-phospho-RPS6$S^{240}$ [1:5000, (9)] and anti-RPS6 antibodies (1:1000, sc-74459, Santa Cruz).

**Immunoblot analyses**

For immunoblotting, proteins were resolved by SDS-PAGE and transferred to PVDF membranes for 90 min 110V at 4°C using transfer buffer (192 mM glycine, 25 mM Tris, 0.1% SDS, 20% ethanol) and a Bio-Rad wet blotting transfer system. Membranes were blocked for at least 1h (5% w/v non-fat dry milk in 1X TBS, 0.05% Tween®) and then incubated with the relevant primary antibody under gentle rocking overnight at 4°C. Secondary antibodies conjugated with horseradish peroxidase (Jackson ImmunoResearch) were used at 1:20000 in 5% non-fat milk in TBS for 1h at RT. Chemiluminescence was performed using a SuperSignal West Femto Maximum Sensitivity Substrate (Thermo Scientific). Images were acquired using ChemiDoc system (Biorad) equipped with a CCD camera.

**Chemicals**

Stocks of ABA (Duchefa Biochemie A0941; 10 mM stock in 50 mM Tris-HCl pH 8.5) and Leptomycin B (Santa Cruz Biotechnology sc-202210; 1 mM stock in ethanol), were prepared and stored at -20°C and used at the indicated concentrations.

**Statistical analysis**

Basic data processing was performed in Excel. Statistical analyses were performed using GraphPad Prism version 8.4.0 for Windows, GraphPad Software, La Jolla California USA.
